## Supplementary files for "Cell-free RNA reveals host and microbial correlates of broadly neutralizing antibody development against HIV"

### Supplementary Figures

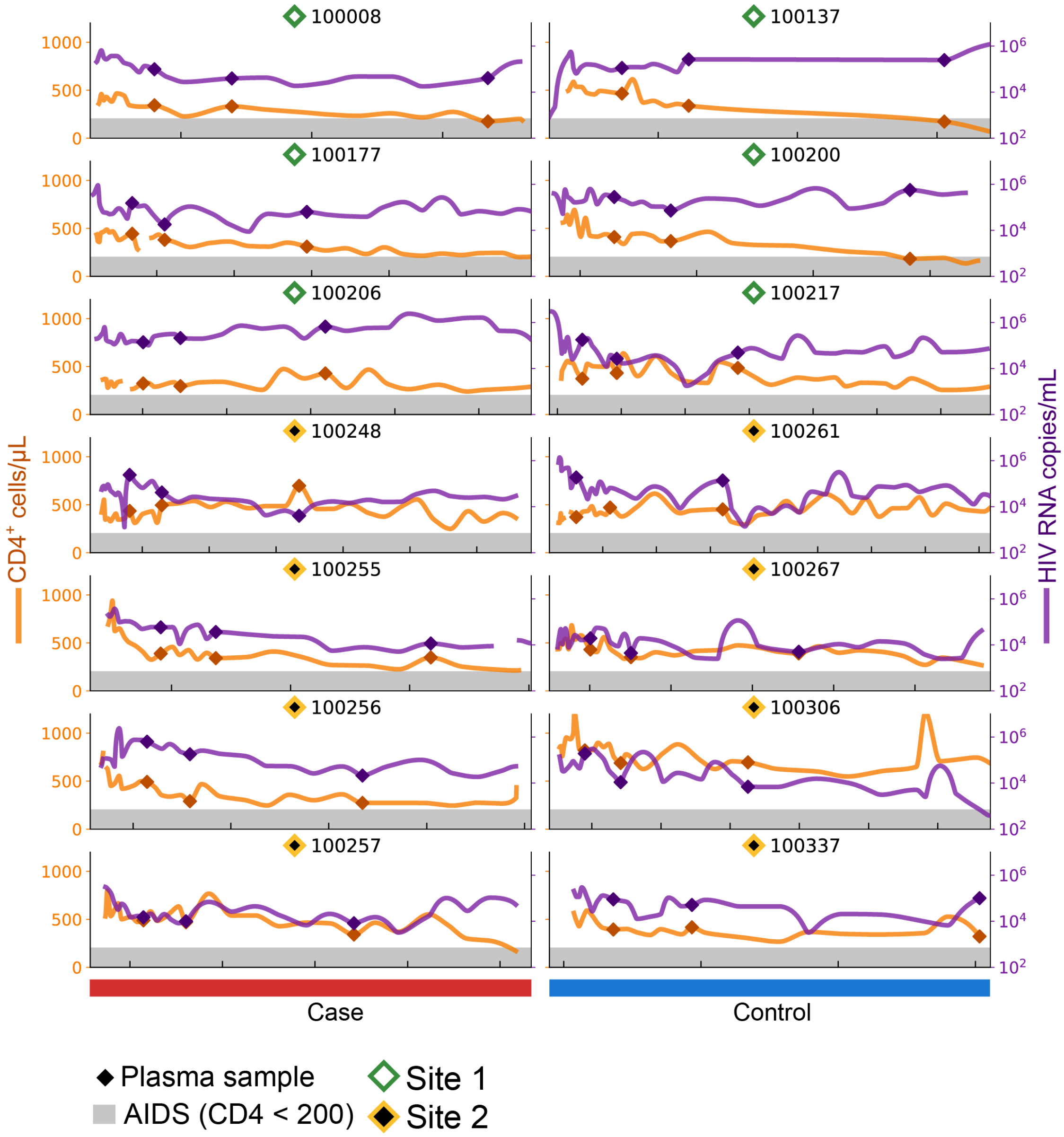


**Figure S1: Clinical information for blood samples.** Longitudinal trajectories for CD4 (orange) and viral load (purple). Diamonds indicate timepoints sequenced in this study. Grey bars indicate values for CD4 count < 200.

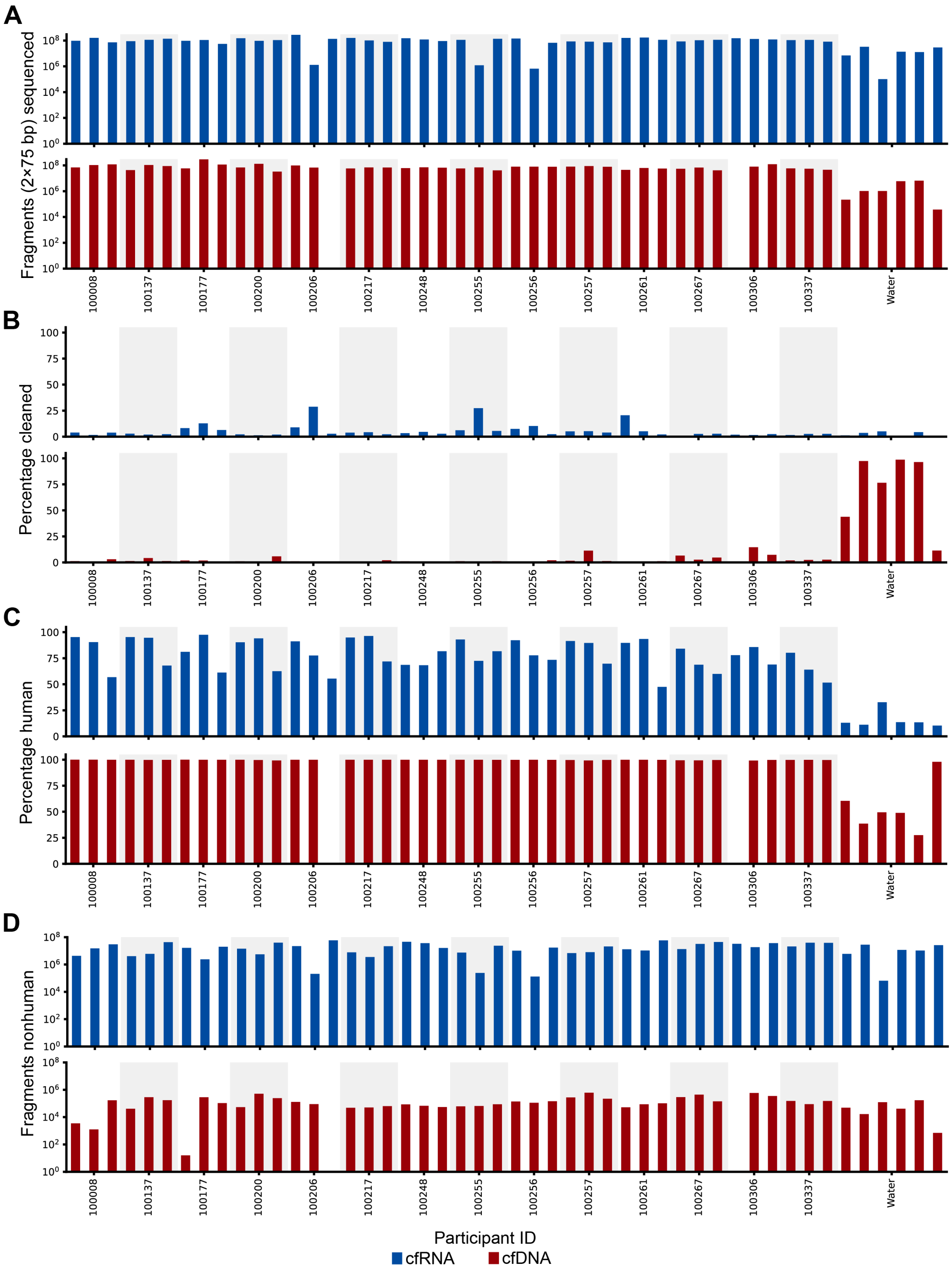


**Figure S2: Sequencing quality control metrics per sample.** A) Number of reads (2x75 bp) sequenced in each sample and negative controls. Two cfDNA samples failed extraction/library preparation. B) Percentage of reads cleaned in QC steps (low quality or that align to the UniVec Core database). Percentages are generally very low, except in cfDNA negative controls, which had a high proportion of primer/library-derived sequences. C) Percentage of reads mapping to the human genome. D) Total number of non-human reads, typically more than a million for the cfRNA samples and around 10 thousand for the cfDNA samples. Samples are ordered by time within each participant.

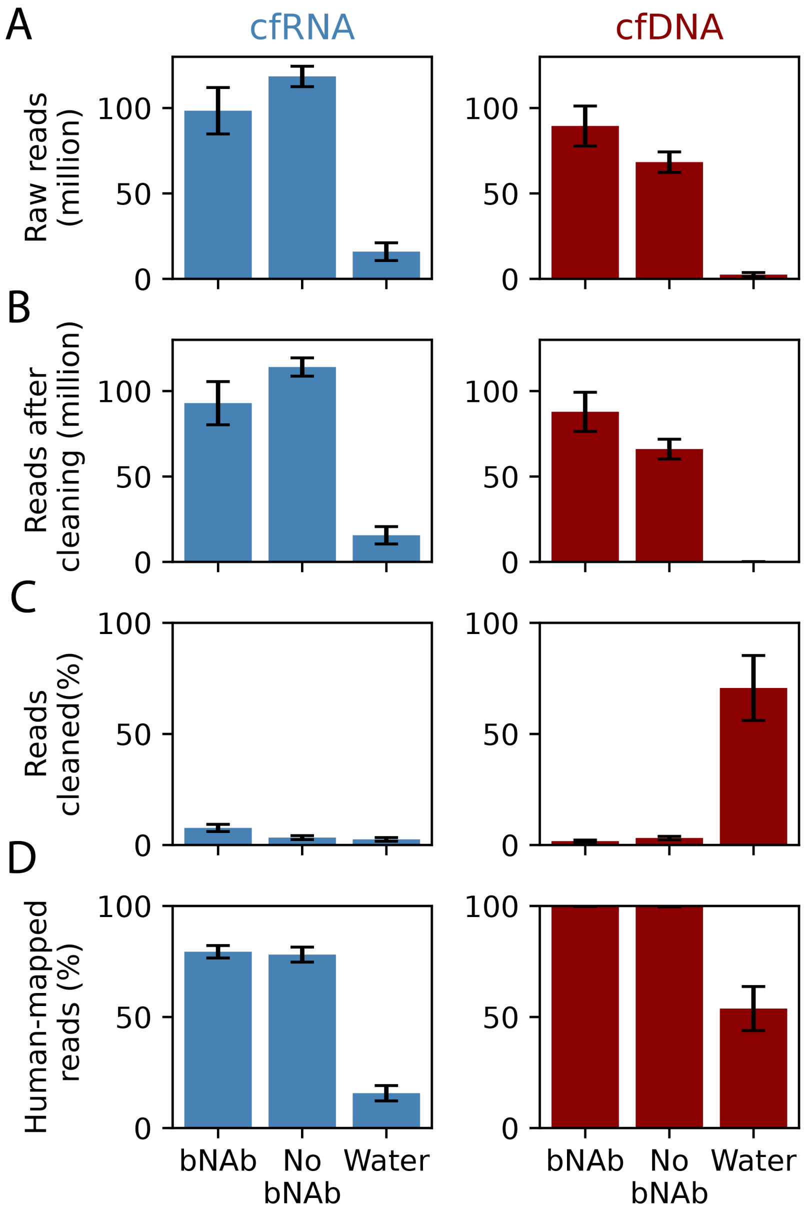


**Figure S3: Sequencing quality control metrics summarized by study group.** A) Total sequencing depth per sample (raw read counts). B) Read counts retained after QC cleaning steps (see Methods) C) Percentage of reads removed during cleaning D) Percentage of human-mapped reads after cleaning. Metrics are shown for cfRNA (left) and cfDNA (right) libraries across bNAb producers, non-producers, and negative controls. Data are shown as mean ± SEM.

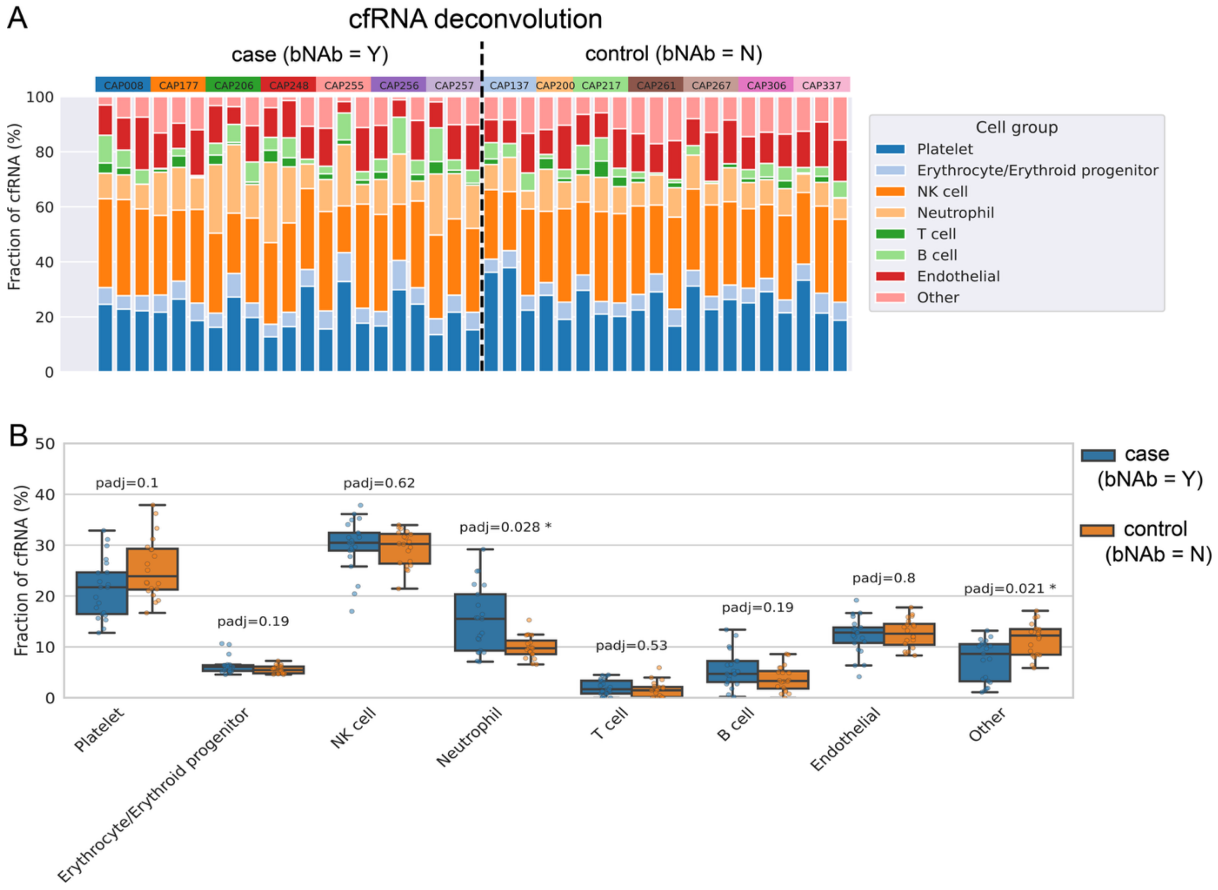


**Figure S4. Cell-type deconvolution analysis of cfRNA.** (a) Relative cell-type contributions to the cfRNA composition for each sample. Samples are grouped by participant and study group (bNAb producer or control), and less abundant cell types are aggregated into 'Other'. (b) Box plots showing the relative abundance of each cell type in bNAb producers and controls. Statistical significance was assessed using a Mann–Whitney U test, with p-values adjusted for multiple hypothesis testing using Benjamini–Hochberg.

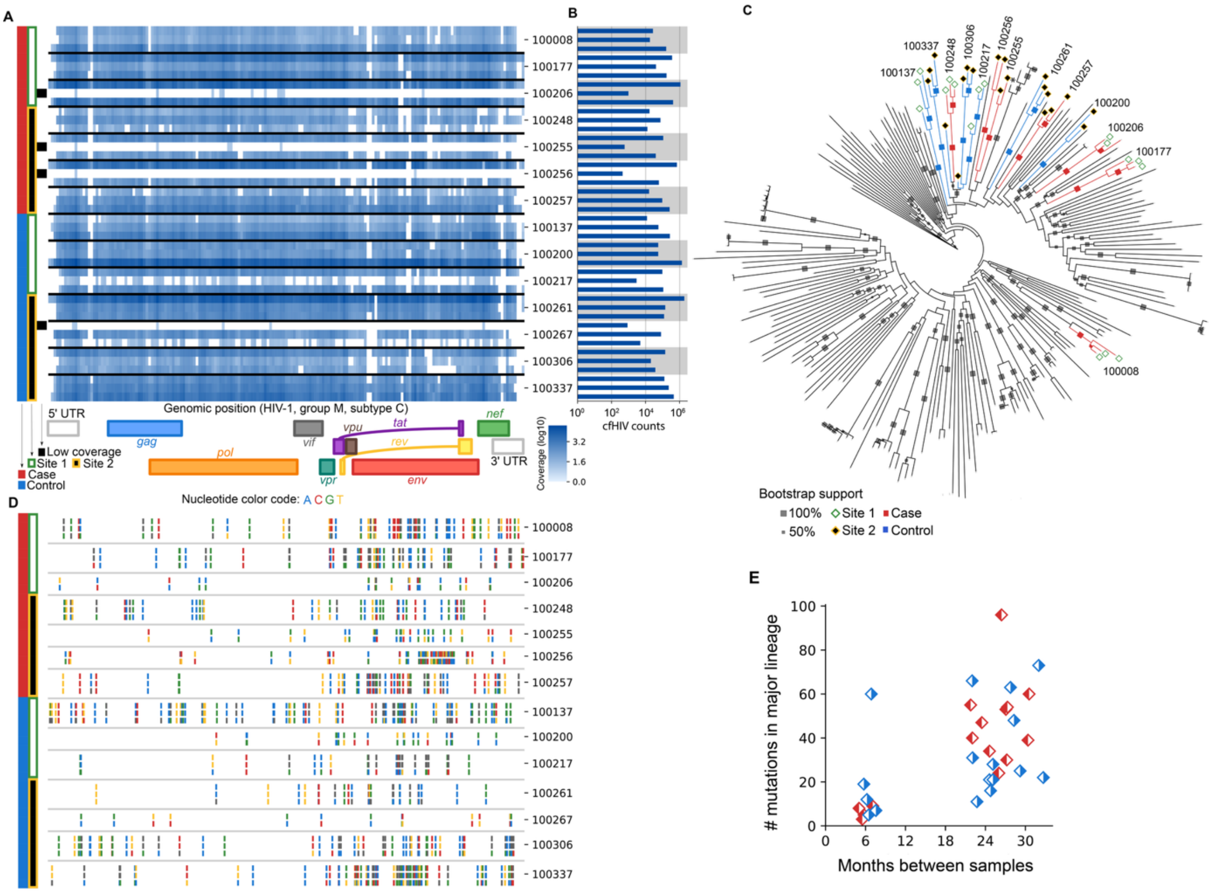


**Figure S5: HIV genotypes, coverage and mutations.** A) Heatmap of coverage across the genome in each sample. The four low abundance samples are marked with black squares on the left hand panel. B) Total number of read fragments aligning to HIV in each sample. C) Phylogenetic tree of HIV consensus genotypes for each sample obtained in this study together with 200 other genotypes of HIV-1 group M, subtype C from South Africa. D) Locations of SNVs local to each participant across the three time points. Most cluster within the *env* gene. E) Mutation rate observed in the samples using differences in the consensus genotypes within participants.

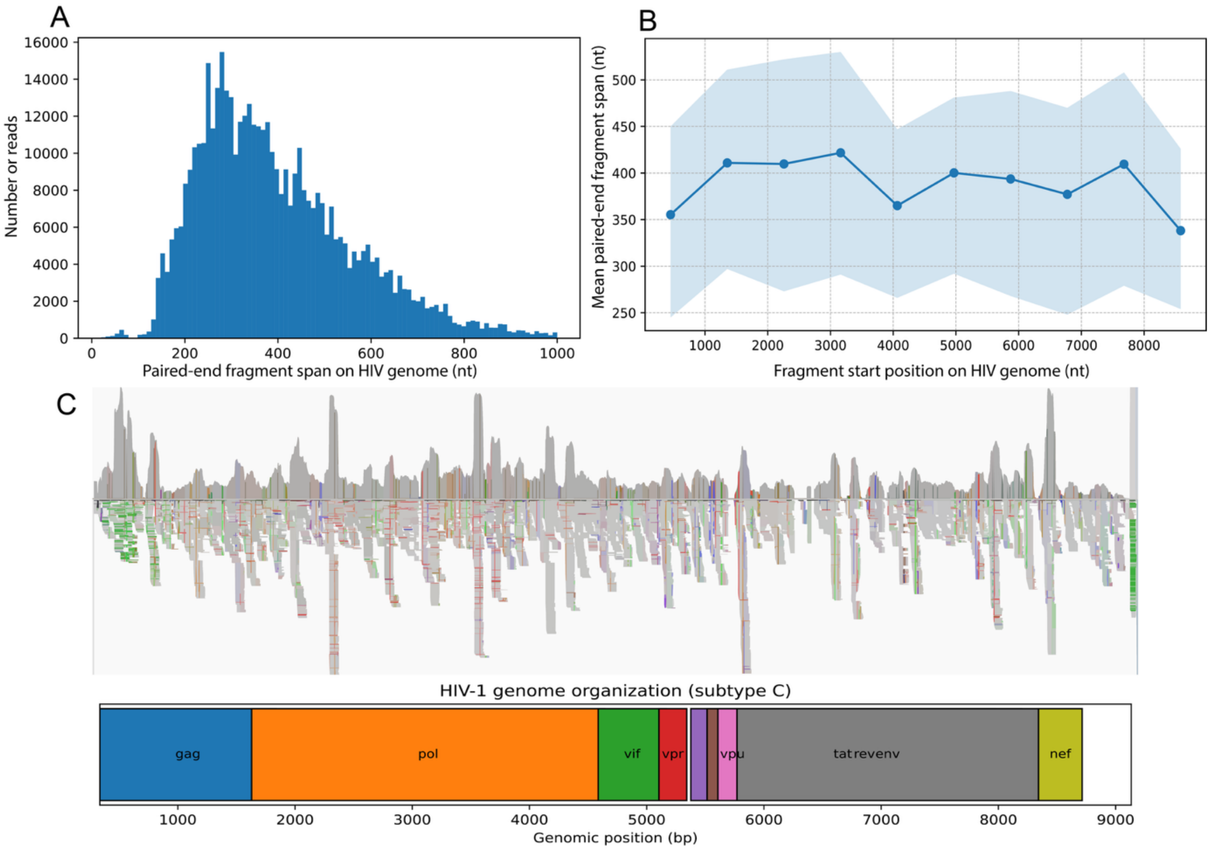


**Figure S6: Fragment length and coverage of cfHIV reads.** A) Distribution of paired-end genomic spans for reads mapping to the HIV genome, showing a predominance of short fragments (200-600 bp). B) Mean paired-end genomic span as a function of genomic start position, showing no systematic trends that could indicate priming from long RNA templates. (c) Genomic coverage profile across the HIV genome, shown alongside the reference organization of the HIV genome.

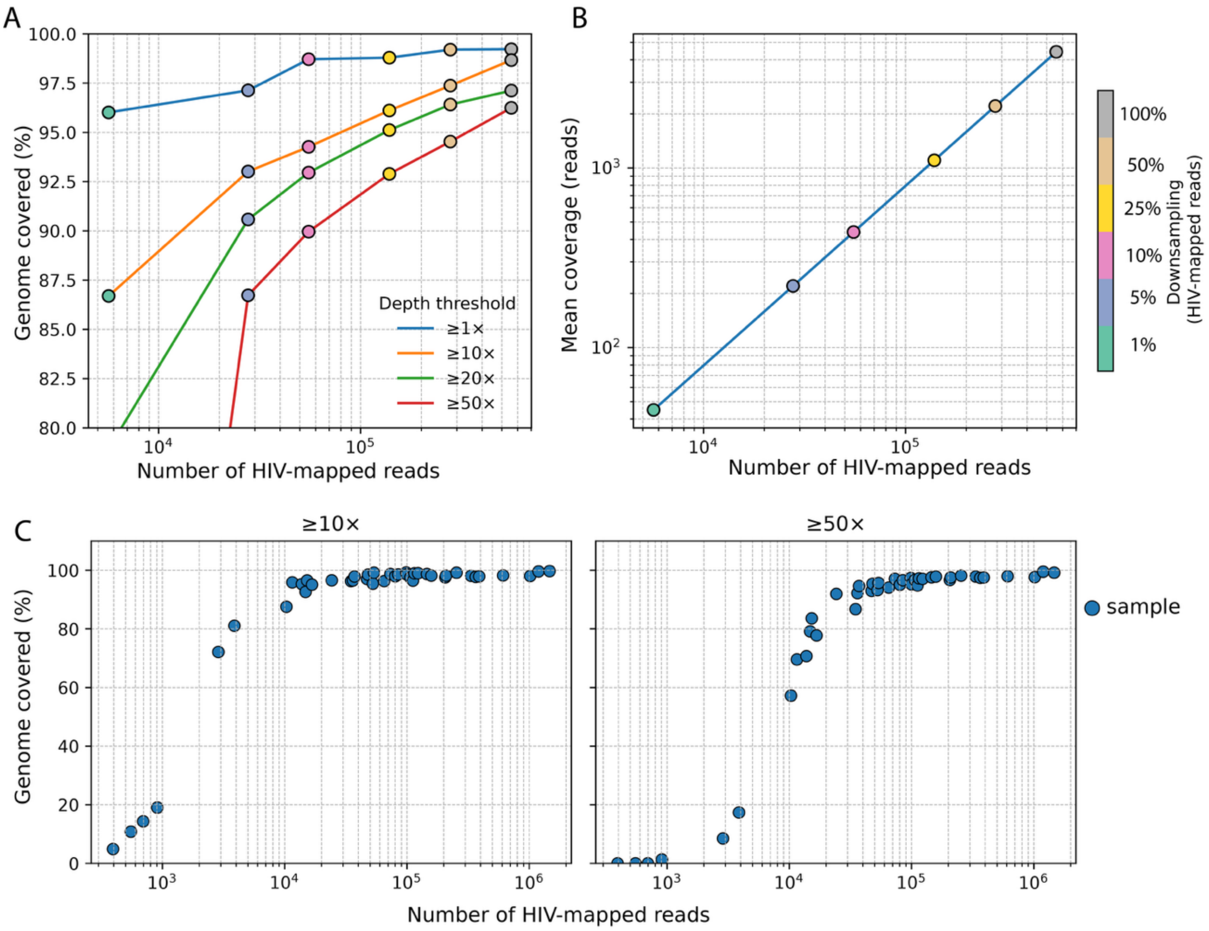


**Figure S7: Subsampling analysis of HIV genome coverage from cfRNA.** A) Fraction of the HIV genome covered above increasing minimum depth thresholds (1X-50X) at different subsampling levels of sample CAP261. B) Number of HIV-mapped reads versus mean coverage depth across the HIV genome, showing a linear relationship. C) Genome coverage achieved for each sample in the cohort at ≥10X (left) and ≥50X (right) depth thresholds.

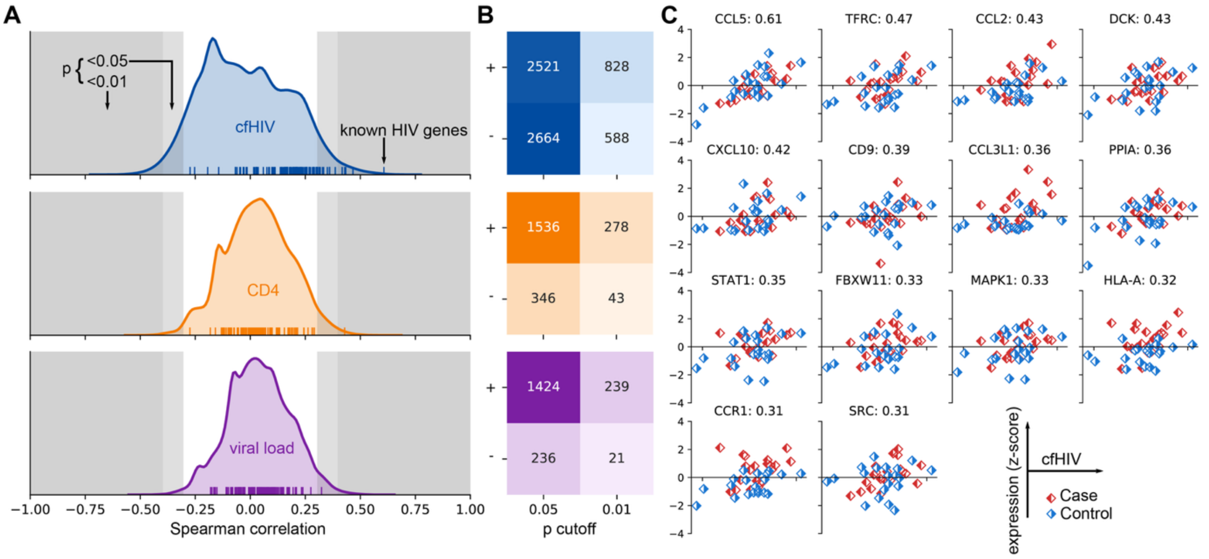


**Figure S8: Correlations of genes with different disease measurements.** A) Distributions of Spearman correlations of all genes with either cfHIV counts (top, blue), CD4 counts (middle, orange), or viral load (bottom, purple). Gray bands indicate regions of significant correlation as determined by the Spearman test on the data. The small ticks (rugplot) at the bottom of each plot indicate known HIV genes. B) Tables of the number of genes strongly correlated/anticorrelated between genes and measurements (as in A). C) Scatter plots of cfHIV abundance vs gene expression (z-score) for the 14 significantly correlated known HIV-associated genes. Besides HLA-A, none of these show strong differences between the two study groups.

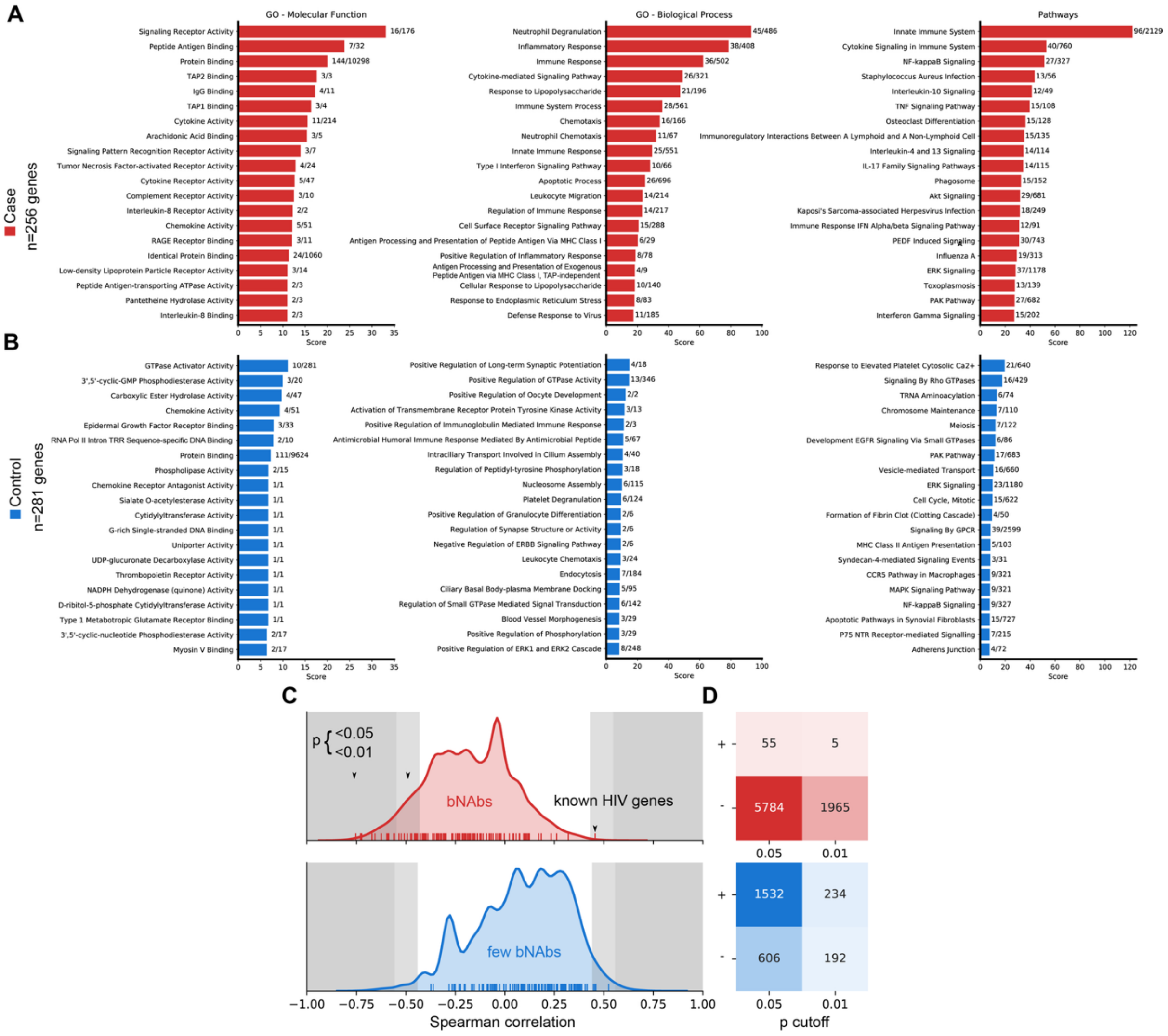


**Figure S9: Enriched gene pathways in bNAb producers.** A) Top gene ontologies (molecular function, biological process) and pathways enriched in genes that are elevated in bNAb producers. Numbers indicate the number of selected genes compared with the total number assigned to that category. B) Similar to A, but for the genes with decreased abundance in bNAb producers. Note that scores are much lower for decreased genes than for elevated genes and no clear enriched groups of genes are present in decreased genes. C) Distribution of genes correlated with breadth in bNAb producers (upper, red) and controls (lower, blue). D) Tables of the number of genes meeting correlation thresholds in (C).

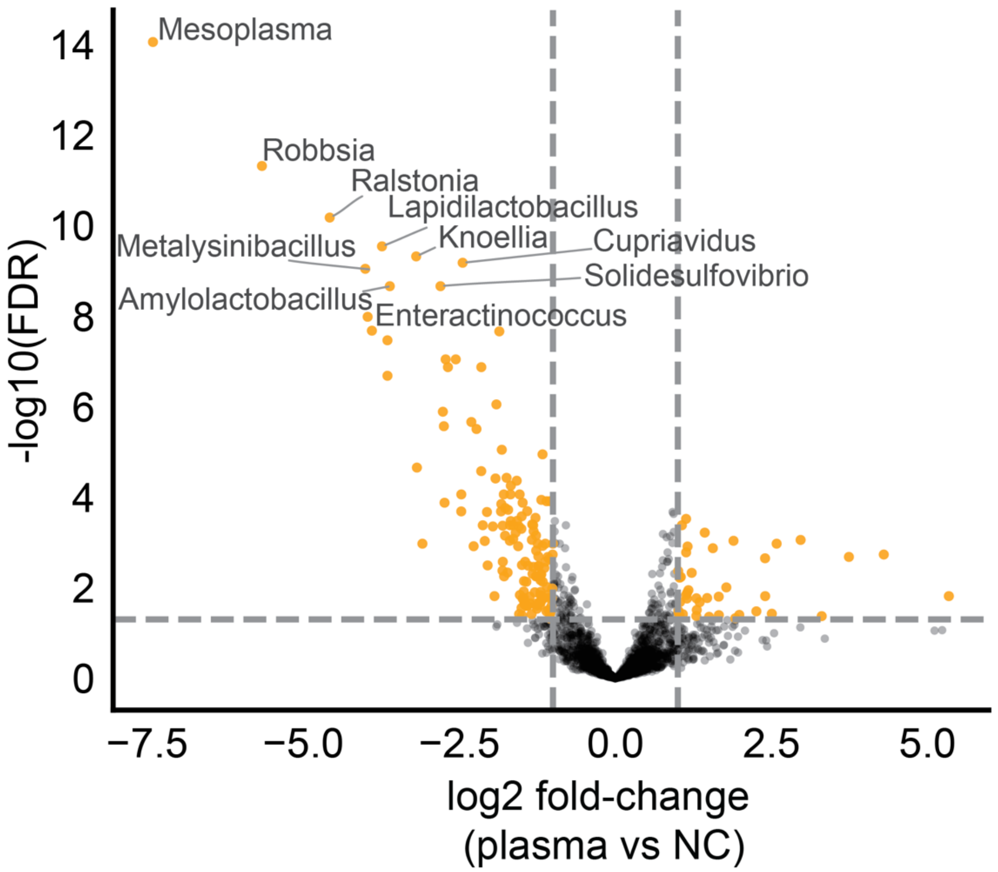


**Figure S10: Background microbial sequences in negative controls.** Volcano plot showing differential abundance of microbial genera between plasma samples and negative controls (NC) (FDR < 0.05, Benjamini–Hochberg). Negative log2 fold-change values indicate genera enriched in NC, which were used to define background levels. Several NC-enriched taxa showed large effect sizes ($|\log_2 FC|>4$, $FDR<10^{-8}$), primarily comprising environmental taxa previously reported in DNA extraction kits and reagents in low-biomass sequencing.

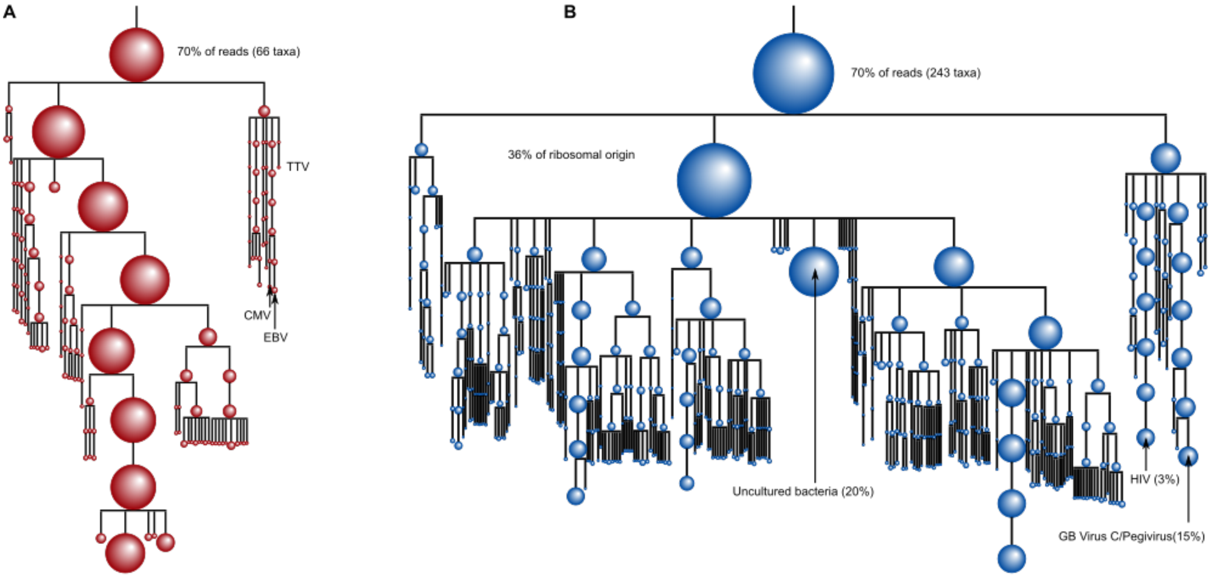


**Figure S11: Phylogenetic tree of microbial sequences from circulating nucleic acids in HIV infection (all participants).** A) Phylogenetic tree of cfDNA-derived microbiome reads pooled across all participants (PLWH who do and do not develop bNAbs). The main branches represent archaea, bacteria and viruses (from left to right); representative viruses including Torque Teno Viruses (TTV), Epstein-Barr virus (EBV) and cytomegalovirus (CMV) are highlighted. B) Phylogenetic tree of cfRNA-derived microbiome reads pooled across all participants. The main branches represent archaea, bacteria and viruses (from left to right). Within the viral branch, HIV and GB virus C are the main contributors to the virome.

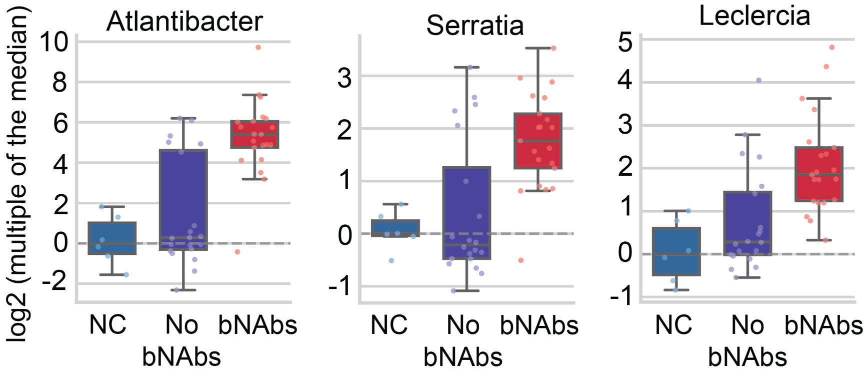


**Figure S12: Subset of microbial genera enriched in bNAb producers.** Boxplots showing normalized abundances of selected microbial genera across negative controls (NC), bNAb non-producers, and bNAb producers. Values are shown as log2-transformed abundances relative to the median abundance in negative controls.

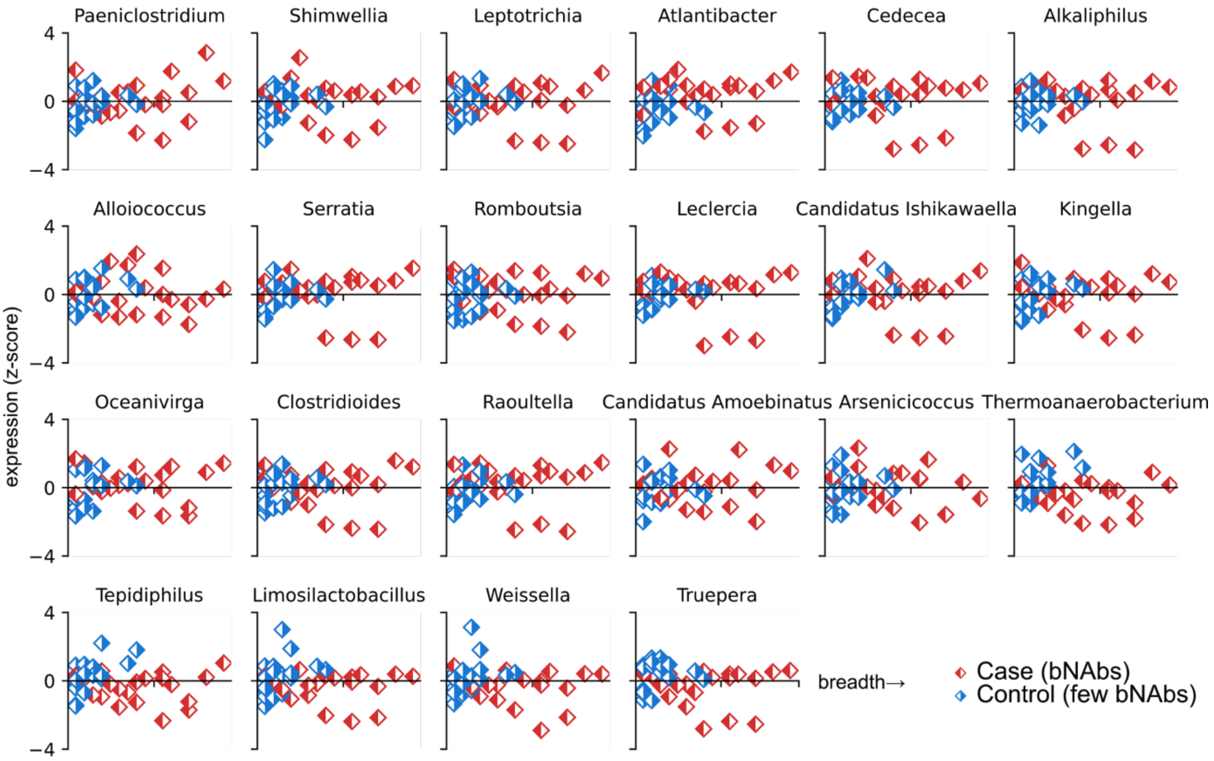


**Figure S13: Pseudo-temporal dynamics in differentially abundant genera between bNAb producers and non-producers.** Scatter plots of the abundance (z-score) vs breadth production (%) for the 22 genera found to be differentially abundant between case (bNAbs) and control (no bNAbs). No apparent time differences found.

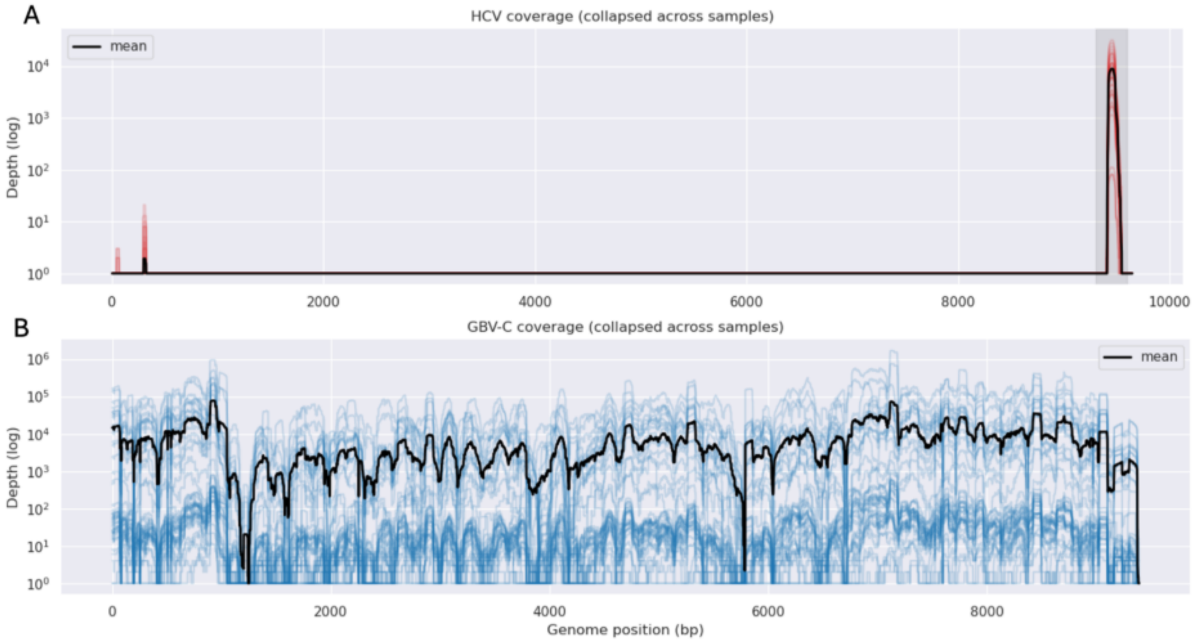


**Figure S14: Coverage profiles of HCV and GBV-C genomes across samples.** A) Read coverage across the HCV genome. B) Read coverage across the GBV-C genome. Coverage for individual samples is shown in red (HCV) and blue (GBV-C); mean coverage is shown in black.

### Supplementary Tables

**Table S1: Excel file**

**Table S2: Participant codes and corresponding accession numbers in the HIV database (https://www.hiv.lanl.gov/content/index) for the reference FASTA sequences included in this study.**

| Participant code | Accession name |
|---|---|
| CAP8 | GQ999972 |
| CAP137 | JF704693 |
| CAP177 | FJ443379 |
| CAP200 | FJ443274 |
| CAP206 | GQ999982 |
| CAP217 | MN097584 |
| CAP248 | GQ999987 |
| CAP255 | GQ999988 |
| CAP256 | KF241776 |
| CAP257 | GQ999990 |
| CAP261 | NA (Sequence ID: CAP261.pl26) |
| CAP306 | KC154019 |
| CAP337 | OQ551953 |
| CAP239 | GQ999991 |
